# The impact of reporter kinetics on the interpretation of data gathered with fluorescent reporters

**DOI:** 10.1101/834895

**Authors:** Bernardo L. Sabatini

## Abstract

Fluorescent reporters of biological functions are used to monitor biochemical events and signals in cells and tissue. For neurobiology, these have been particularly useful for monitoring signals in the brains of behaving animals. In order to enhance signal-to-noise, fluorescent reporters typically have kinetics that are slower than that of the underlying biological process. This low-pass filtering by the reporter renders the fluorescence transient a leaking integrated version of the biological signal. Here I discuss the effects that low-pass filtering, or more precisely of integrating by convolving with an exponentially decaying kernel, has on the interpretation of the relationship between the reporter fluorescence transient and the events that underlie it. Unfortunately, when the biological events being monitored are impulse-like, such as the firing of an action potential or the release of neurotransmitter, filtering greatly reduces the maximum correlation coefficient that can be found between the events and the fluorescence signal. This can erroneously support the conclusion that the fluorescence transient and the biological signal that it reports are only weakly related. Furthermore, when examining the encoding of behavioral state variables by nervous system, filtering by the reporter kinetics will favor the interpretation that fluorescence transients encode integrals of measured variables as opposed to the variables themselves. For these reasons, it is necessary to take into account the filtering effects of the indicator by deconvolving with the convolution kernel and recovering the underlying biological events before making conclusions about what is encoded in the signals emitted by fluorescent reporters.

## Introduction

Genetically-encoded fluorescent reporters of biological processes are revolutionizing neurobiological and cell biological research. Specifically within neurobiological research, reporters of intracellular calcium (e.g. GCAMPs, RCAMPs, …) are used as surrogate measures of action potential firing whereas reporters of neurotransmitters and neuromodulators (e.g. GluSFNRS, dLight, and DA-GRAB, …) are used to detect the release of such molecules (Akerboom et al., 2013; Akerboom et al., 2012; Marvin et al., 2018; Patriarchi et al., 2018; Sun et al., 2018).

Fluorescent reporters generally consist of genetically-encoded fluorophores into which a ligand binding site has been engineered that, when occupied, alters the fluorescence properties of the chromophore. The kinetics of the observed fluorescence transient depend on many factors, including the time varying concentration of the ligand, the on and off rates of ligand binding to the reporter, and the kinetics of induction and reversal of the conformational change in the reporter downstream of ligand binding and unbinding. The kinetics of ligand binding and downstream conformational changes typically make the kinetics of the fluorescence transients a low-pass filtered version of the kinetics of the ligand concentration.

These filtering effects can range from potentially inconsequential to severe. On the fast side, synaptically released glutamate stays high in the synaptic cleft for < 1 ms (Clements et al., 1992) and triggers ∼25 ms fluorescence transients for the newest versions of glutamate reporters (Durst et al., 2019; Marvin et al., 2018) which, if an individual synapse is active at less than 40 Hz, may not lead to accumulation of signal across release events. On the slow side, a ∼1 ms long action potential and the calcium current that it generates results in long-lasting changes in intracellular calcium concentration and GCAMP fluorescence that last ∼1-10 seconds. In general, from the perspective of signal-to-noise optimization, it is advantageous to make the kinetics of the reporter as slow as is biologically acceptable, as this allows more photons to be collected and analyzed to judge if an event, such as a action potential or vesicle fusion, has occurred.

Although the slowness of fluorescent reporters is advantageous for signal detection, it can be a problem when trying to decipher what is encoded by activity in the nervous system. Many studies attempt to relate action potential firing or neuromodulator release to behavioral events in order to understand what is encoded by the activity of neurons or synapses. In this case it is often necessary to compare impulse or impulse-like events (e.g. playing a tone, generating a light flash, licking a water spot, receiving a reward…) to the slow fluorescence signals generated by the reporters described above. Other studies attempt to relate neural activity to more abstract variables such as motivation and value that are calculated from animal behavior. In both cases, the general approach is to infer what is encoded by the biological process reported by the sensor by calculating the correlation or covariance between the time courses of fluorescence and behavioral events.

Here I discuss why it is necessary to consider the kinetics of the fluorescent reporter when studying the relationship between biological and behavioral events. I show that even mild temporal filtering by the fluorescent reporter severely dampens the maximum correlation coefficient that can be observed between fluorescence and the events that underlie the fluorescence changes. Furthermore, I show that these effects make the interpretation of the relationship between fluorescence transients and biological or behavioral events difficult and can lead to erroneous conclusions. Lastly, I discuss methods to deal with these challenges.

## Results

### Modeling fluorescence transients as convolutions

Consider an event that occurs at t=0 sec that induces an increase in fluorescence from a reporter (Figure 1). Here I approximate the event as being an impulse (i.e. ideally last no time but in practice assigned to a single time bin) and the fluorescence transient as rising instantaneously and decaying exponentially with a single time constant (Figure 1a). This fluorescence transient that results from an impulse-like event is referred to as the impulse response function and can be used as a convolution kernel to generate the fluorescence transients expected from the linear summation of many impulse-like events (Figure 1B).

**Figure 1.**
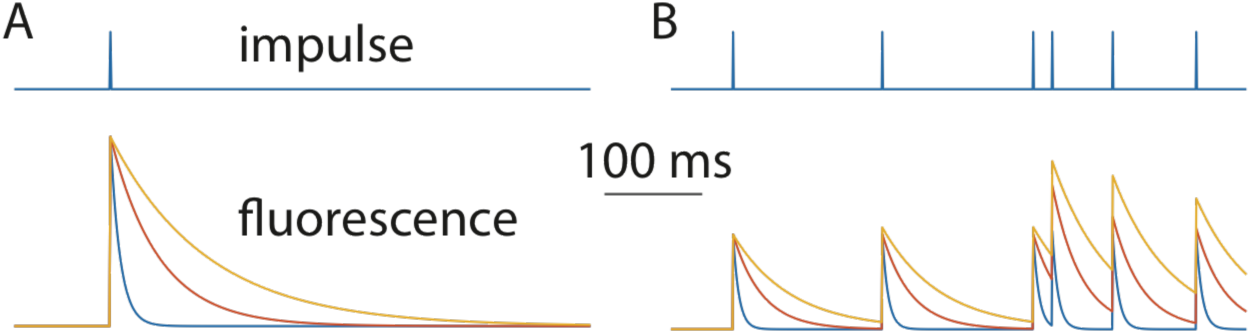
Filtering of impulse-like events by the kinetics of fluorescent reporters. **A**, A single impulse (top), such as an action potential, can elicit long-lived fluorescent transients (bottom) whose kinetics are determine by the decay time constant of an exponential kernel. Data are shown for kernels with 10, 50, or 100 ms time constants (blue, red, orange). **B**, As in panel A for a random series of impulses (top) leading to summation of fluorescent signals (bottom) with the degree of summation depending on the time constant of decay of the convolution kernel.

### The correlations between fluorescence transients and the events that create them

In the above example the fluorescence transient is completely and solely determined by the timing of the impulses. However, because the convolution kernel spreads the impulse into neighboring time bins, the correlation between the impulse and the modeled fluorescent transient is not perfect: the correlation coefficient between the impulse and the modeled fluorescent transient will be less than one and is capped at a maximum value determined by the impulse response function. For the examples shown in Figure 1, the correlation coefficients between the impulse-like biological events and the fluorescence transients they generate fall to 0.43, 0.20, and 0.14 for exponential convolution kernels with time constants of 10, 40, and 100 ms, respectively. The results are similar when considering a single event (Figure 1A) or a random series of events (Figure 1B). Note that for clarity of presentation I have chosen to use time units of ms but, of course, the results hold with any units and the analysis can be carried out dimensionless as well.

This result holds for a wide range of time constants of the exponential convolution kernel (Figure 2). The maximal correlation coefficients of a train of impulse-like events and the filtered version of these events (Figure 2A) is less than 1 and falls rapidly for time constants longer than the sampling interval. For events at a rate of 10 Hz and assigned to 1 ms time bins (Figure 2B) the maximal correlation coefficient falls below 0.5 even with a 7 ms kernel decay time constant.

**Figure 2.**
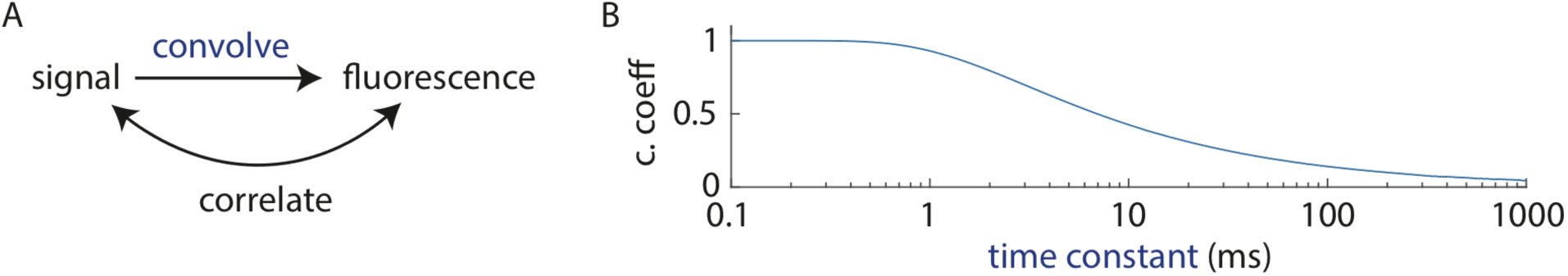
The kinetics of the fluorescent reporter limit the maximal correlation coefficient that can be observed between fluorescence and the underlying impulse-like biological process. **A**, Schematic of the analysis. An impulse-like biological signal is convolved with an exponential kernel to mimic the effects of the kinetics of the fluorescent reporter. The cross-correlation between the ensuing fluorescence and the original signal is calculated. **B**, The correlation coefficient of an underlying process and the modeled fluorescent transient (as in A) for exponential convolution kernels with varying time constants (blue), ranging from 0.1 to 1000 ms. The signal was modeled as impulse like events each lasting 1 ms and occurring at a rate of 10 Hz. All kernels with decay time constants significantly longer than the time bin duration (1 ms) greatly reduce the maximal detectable correlation coefficient.

Given the ∼30 Hz frame rate of many cameras and laser-scanning microscopes, a sensor with 100 ms kinetics is about as fast as one would like as it forces the estimate of amplitude from ∼3 samples. Nevertheless, these results indicate that, even under optimal conditions (i.e. no noise or time lags), the maximal correlation coefficient that can be expected between the fluorescence of such a sensor and the impulse-like events that generate the fluorescence changes is very small (0.14).

How can these effects impact the interpretation of fluorescence transients? Consider events occurring at a rate of 10/sec and once again filtered by an exponential kernel with a decay time constant of 100 ms. This approximates the case of attempting to relate dopaminergic neuron action potential firing (basal rate of 5-10 Hz) measured with an electrode to dopamine concentrations measured with dLight (Patriarchi et al., 2018), a sensitive and bright fluorescent dopamine reporter with an impulse response on the order of 100 ms. As described above, even if the dopamine concentration is solely and completely determined by the activity of the neurons recorded, the maximal correlation coefficient between the spikes and the fluorescence emitted by the dopamine sensor will be only 0.14. Such a low correlation coefficient might erroneously lead to the conclusion that most of the variance in dopamine concentration is not due to the variable firing of dopamine neurons but is instead caused by another factor.

### Correlations between fluorescence transients and behavioral variables

Failing to find a correlation between dopamine neuron spiking and dopamine levels, one might try to determine what else controls dopamine levels and examine the behavioral variables that are encoded by the dopamine transients, as reported by dLight fluorescence. Variables derived from behavioral monitoring fall into two general categories. First, there are instantaneous variables that can be directly observed or related to ongoing behavior and occur at precise times. These include the onset of cues used in operant or Pavlovian conditioning (e.g. light flashes or tones) or simple motor actions (e.g. licks or arm movements) used either to indicate choice or collect reward. Similarly, reward-prediction errors are instantaneous variables as they are calculated at and assigned to the moment when the animal discovers the presence or absence of a reward. Second, slowly-changing behavioral variables can be calculated from the leaky integration of behavioral events. These are often referred to as “state” variables. For example, motivation, value, and attention, might be calculated from integrated measures of movement latencies, reward history, and salient events, respectively. Leaky integration is mathematically equivalent to performing a convolution, with the time constant of the kernel reflecting the time over which information is retained (i.e. how quickly it leaks out).

Analysis of the encoding of the first class of behavioral variables by the fluorescence emmitted from a reporter is confounded by the same considerations presented above. The ability to relate instantaneous behavioral variables or events to fluorescent signals that reflect low-pass filtered neuronal events is limited by the time constant of filtering of the reporter with correlation coefficients capped as in Figure 2.

The situation is worse when considering the second class of behavioral variables (Figure 3). An investigator will typically perform a leaky integration over behavioral events to calculate a behavioral state variable hypothesized to explain the fluorescence transients (Figure 3A). The investigator may then examine if the observed fluorescent transient is best correlated to the underlying impulse-like behavioral events or to the calculated state variables. For example, one might ask if dopamine transients reported by dLight are better correlated with either (1) instantaneous reward prediction error (RPE) assigned to the moment of reward presentation or omission or (2) an expected value of the motor action that is calculated based on the history of previous action outcomes and timing.

**Figure 3.**
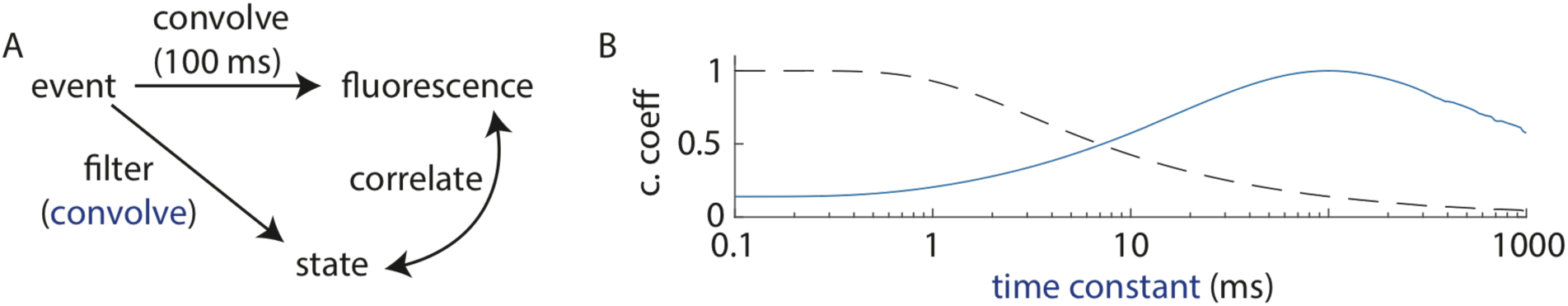
The kinetics of the fluorescent reporter enhance the correlation coefficient between the observed fluorescent transient and low-pass filtered versions of the underlying events. **A**, Schematic of the analysis. A series of impulse-like biological events is convolved with an exponential kernel (100 ms time constant in this case) to model the effects of the slow kinetics of a fluorescent reporter and generate a fluorescence transient. The event series is also used to calculate a “state” behavioral variable with an exponential filter (blue) of varying kinetics. The cross-correlation between the fluorescence transients and these state variables are calculated. **B**, Using the analysis from panel A, the correlation coefficients between fluorescent transients (modeled with fixed 100 ms time constant) and state variables are calculated as a function of the time constant (0.1-1000 ms) of the exponential convolution kernels (blue) used to generate the state variable. The event series was modeled as impulse like events each lasting 1 ms and at arising at rate of 10 Hz. The correlation coefficient if high (> 0.5) for a wide range of convolution kernels with time constants from ∼10-1000 ms. The data from 2B are replotted in the dashed line for comparison.

Here we consider the case in which the same sequence of events generates both the fluorescence transient and the behavioral state variable – i.e. the fluorescence transient and the state variable are completely and solely linearly determined by the same impulse train (Figure 3A). An example is the case in which modulation of dopamine neuron firing is linearly related to RPE, dopamine levels are measured with dLight, and the experimenter asks if the dLight transient is better explained by RPE or value as calculated by leaky integration of RPE.

Figure 3B show the correlation coefficients between (1) the fluorescence generated by convolving impulse events with an exponential kernel with a 100 ms time constant and (2) a “state” variable calculated from convolving the impulse events with exponential decay kernels of varying time constants. Of course, when the time constant of leaky integration used to calculate the state variable is also 100 ms, the correlation coefficient is maximal and equal to 1. However and surprisingly, a correlation coefficient above 0.5 is found for any time constant in the range of ∼9-800 ms (Figure 3B). Thus, even over very broad definitions of the state variable, the correlation coefficient of the fluorescence transient to the state variable is high. Furthermore, it appears high especially when compared to the low coefficient calculated between the fluorescence transient and the impulse events used to generate both signals (Figure 3B).

These effects are robust and essentially constant over a broad range of impulse event rates and convolution kernels time constants (Figure 4). The 100 ms filtered calculated fluorescence transient remains poorly correlated to the underlying impulse-like events (Figure 4A) and relatively well-correlated to calculated state variables (Figure 4B) over 3 and 4 orders of magnitude changes in the impulse rates and leaky integrator time constants, respectively.

**Figure 4.**
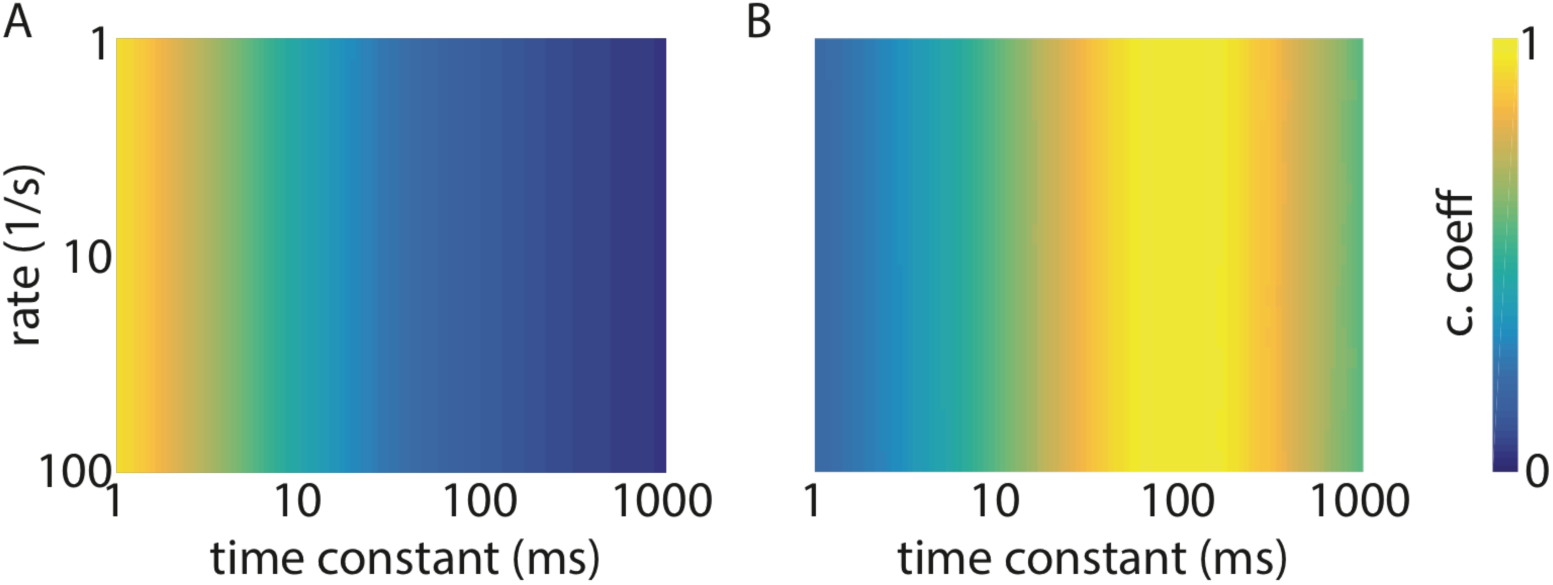
Robustness of the effects shown in Figure 2 and 3. **A**, Color coded correlation coefficients (blue to yellow, 0 to 1) between the modeled fluorescence transients and the impulse-like events that generated it (as in Figure 2) for a range of event rates (y-axis) and convolution kernel decay time constants (x-axis). **B**, Color coded correlation coefficients (blue to yellow, 0 to 1) between the modeled fluorescence transient (fixed 100 ms decay) and a calculated state variable (as in Figure 3) for a range of event rates (y-axis) and state variable leaky time constants (x-axis).

## Discussion

Many neurobiologists strive to understand how animals integrate information to make decisions and thus exploit fluorescent reporters to reveal the features of the environment and behavioral history that are encoded in the nervous system. This manuscript is not meant to denigrate such efforts or to argue against the utility of fluorescent reporters in such efforts. On the contrary, it is meant to help such efforts by pointing out potentials pitfalls, which, luckily, can be avoided.

Fluorescent reporters are typical designed to optimize signal detection, which favors slow reporters that increase signal-to-noise by extending the fluorescent signal well beyond the lifetime of fast underlying events. However, the kinetics of useful reporters are also bounded by the dynamics of the signal of interest, which favors fast reporters whose signals decay to baseline on a time scale similar to the inter-event interval. This balance typically results in useful reporters being those whose fluorescence reflects a leaky integration of one to tens of underlying events (e.g. ∼1 s decay of a genetically-encoded calcium indicator used to monitor ∼10 Hz neuronal firing).

Fluorescence reporters with kinetics optimized for signal detection become problematic when used to calculate correlations to impulse-like behavioral events or even to the impulse-like neuronal events that directly generate the fluorescence transients. Since an impulse, by definition, has very limited duration, the correlation of an impulse with its low-pass filtered self is low. Intuitively this makes sense as low-pass filtering takes signal from the impulses and puts it in other time bins, most of which did not have impulses in them. This effect destroys correlation coefficients quickly, which may lead to the erroneous conclusion that a fluorescence transient is not well explained by the series of impulse-like events that generated it.

Similar concerns apply to attempts to use fluorescence transients to discover the behavioral variables that are encoded in the nervous system. Particularly problematic is the analysis of slow time-varying state variables such as value, motivation, and attention, which are typically calculated from the leaking integration of precisely-timed behavioral or sensory events such as the acquisition of rewards, latency or timing of motor actions, and presentation of sensory stimuli. The filtered fluorescence transient will be better correlated with these calculated events than with the impulse events used to generate both. Unfortunately, the good correlation between the two filtered signals holds over a wide range of parameters, which sadly would give the investigator confidence that the conclusions are not very sensitive to the model parameters and thus more likely to be true.

As a the investigator of a laboratory that studies the basal ganglia, brain structures intimately involved in reward reinforcement learning, I am particularly concerned about recent conclusions that the activity of dopaminergic neurons does not explain the concentration of dopamine in their target regions and that the concentration of dopamine is more correlated to the value of an action than to the reward-prediction error generated by the action (Hamid et al., 2016; Mohebi et al., 2019). Although many pieces of data support these conclusions, they were at least in part reached based on the interpretation of data that potentially suffer from the effects described above. Although I phrased the discussion above in terms of fluorescent reporters, the conclusions hold equally well for any slow measurement system, including monitoring dopamine with fast scan cyclic voltammetry which, as traditionally used, produces a signal lasting ∼2 s for a brief optogenetic stimulation of dopamine release.

### Work around

In the linear regime described above, one can correct for the effects of the kinetics of the fluorescence reporter. This can be accomplished by purely analytical approaches such as deconvolution or by fitting algorithms such as constrained iterative deconvolution or modeling with auto-regressive processes. The latter is built into many modern packages for the analysis of fluorescence transients collected from neurons expressing calcium sensitive fluorophores (Giovannucci et al., 2019; Pnevmatikakis et al., 2016; Zhou et al., 2018). Such approaches allow one to extract the activity pattern that best explains the observed fluorescence transient and thus, to an approximation, recover the impulse-like biological events that generate the fluorescence transient and may be related to behavioral and environmental events. An attractive approach is to use a generalized linear model to uncover the impulse response function, the sequence of impulses, and their relationship to behavioral variables (Minderer et al., 2019; Parker et al., 2016). These methods should also be applied when considering fluorescence transients generated by reporters of signals other than calcium and neurotransmitters as well as for both cellular level (e.g. laser scanning and miniscope imaging) and bulk (e.g. fiber photometry) data.

The impulse response of a reporter can be directly measured. For example, an action potential can be triggered in a GCAMP-expressing neuron using electrophysiology or optogenetics while measuring fluorescence (e.g. (Dana et al., 2019)). This will give the kernel needed to convolve the action potential train to model the fluorescence transient or, equivalently, to deconvolve the fluorescence transient to obtain the underlying action potential train (I am setting aside here potential nonlinearities of the fluorescence response). However, the action potential-evoked fluorescence transient is the impulse response of the system (i.e. from action potential to fluorescence) and not the impulse response of the sensor (i.e. from calcium to fluorescence). The former depends on may factors, included biological ones such as the concentration of calcium binding proteins and the activity of calcium pumps in the cell, whereas the latter is a biophysical property of the indicator that determines the kinetics of GCAMP fluorescence following an impulse in calcium concentration.

This distinction between the system and sensor impulse response can be important and which to use for deconvolution depends on the question being asked. For example, the sensor impulse response intrinsic to calcium interacting with GCAMP is relevant if one is studying calcium handling but not if, as is typical, one is using GCAMP as surrogate measure of neuronal spiking. Similarly, if one wants examine if dopamine *concentration* is explained by the firing of action potentials in dopaminergic neurons, then it is sufficient to measure the system impulse response by, for example, optogenetically or electrically activating dopaminergic axons once and measuring the dLight response (as in (Patriarchi et al., 2018)). A similar approach is appropriate to examine if dopamine *release* events track reward-prediction errors or other phasic behavioral events. However, if one wants to ask if dopamine *concentration* is correlated with behavioral events, then it is necessary to use the intrinsic dLight impulse response to dopamine binding and unbinding. This would require measuring dLight fluorescence following an impulse in dopamine concentration, which is difficult to deliver but can potentially be done by uncaging dopamine. Alternatively, the response of dLight to near-instantaneous steps increases or decreases in dopamine concentration delivered with stopped-flow cells can be used to measure on and off rates of dopamine binding to dLight and subsequently derive the impulse response.

### Conclusion

Fluorescent reporters are useful and a mainstay of neurobiological research, including in my lab. Paying attention to the issues raised here and employing the methods described above will ensure that the conclusions drawn from the analysis of fluorescence transients are well founded.

## Methods

Models were run in Matlab using scripts available on Github using the files Figure1.m, Figures2and3.m and Figure4.m (https://github.com/bernardosabatini/impulseCorrelations).

## Acknowledgements

I thank members of the Sabatini lab for helpful discussion. BS is supported by the Howard Hughes Medical Institute.

## References

Akerboom, J., Carreras Calderon, N., Tian, L., Wabnig, S., Prigge, M., Tolo, J., Gordus, A., Orger, M.B., Severi, K.E., Macklin, J.J., et al. (2013). Genetically encoded calcium indicators for multi-color neural activity imaging and combination with optogenetics. Front Mol Neurosci 6, 2.

Akerboom, J., Chen, T.W., Wardill, T.J., Tian, L., Marvin, J.S., Mutlu, S., Calderon, N.C., Esposti, F., Borghuis, B.G., Sun, X.R., et al. (2012). Optimization of a GCaMP calcium indicator for neural activity imaging. J Neurosci 32, 13819–13840.

Clements, J.D., Lester, R.A., Tong, G., Jahr, C.E., and Westbrook, G.L. (1992). The time course of glutamate in the synaptic cleft. Science 258, 1498–1501.

Dana, H., Sun, Y., Mohar, B., Hulse, B.K., Kerlin, A.M., Hasseman, J.P., Tsegaye, G., Tsang, A., Wong, A., Patel, R., et al. (2019). High-performance calcium sensors for imaging activity in neuronal populations and microcompartments. Nat Methods 16, 649–657.

Durst, C.D., Wiegert, J.S., Helassa, N., Kerruth, S., Coates, C., Schulze, C., Geeves, M.A., Torok, K., and Oertner, T.G. (2019). High-speed imaging of glutamate release with genetically encoded sensors. Nat Protoc 14, 1401–1424.

Giovannucci, A., Friedrich, J., Gunn, P., Kalfon, J., Brown, B.L., Koay, S.A., Taxidis, J., Najafi, F., Gauthier, J.L., Zhou, P., et al. (2019). CaImAn an open source tool for scalable calcium imaging data analysis. Elife 8.

Hamid, A.A., Pettibone, J.R., Mabrouk, O.S., Hetrick, V.L., Schmidt, R., Vander Weele, C.M., Kennedy, R.T., Aragona, B.J., and Berke, J.D. (2016). Mesolimbic dopamine signals the value of work. Nat Neurosci 19, 117–126.

Marvin, J.S., Scholl, B., Wilson, D.E., Podgorski, K., Kazemipour, A., Muller, J.A., Schoch, S., Quiroz, F.J.U., Rebola, N., Bao, H., et al. (2018). Stability, affinity, and chromatic variants of the glutamate sensor iGluSnFR. Nat Methods 15, 936–939.

Minderer, M., Brown, K.D., and Harvey, C.D. (2019). The Spatial Structure of Neural Encoding in Mouse Posterior Cortex during Navigation. Neuron 102, 232–248 e211.

Mohebi, A., Pettibone, J.R., Hamid, A.A., Wong, J.T., Vinson, L.T., Patriarchi, T., Tian, L., Kennedy, R.T., and Berke, J.D. (2019). Dissociable dopamine dynamics for learning and motivation. Nature 570, 65–70.

Parker, N.F., Cameron, C.M., Taliaferro, J.P., Lee, J., Choi, J.Y., Davidson, T.J., Daw, N.D., and Witten, I.B. (2016). Reward and choice encoding in terminals of midbrain dopamine neurons depends on striatal target. Nat Neurosci 19, 845–854.

Patriarchi, T., Cho, J.R., Merten, K., Howe, M.W., Marley, A., Xiong, W.H., Folk, R.W., Broussard, G.J., Liang, R., Jang, M.J., et al. (2018). Ultrafast neuronal imaging of dopamine dynamics with designed genetically encoded sensors. Science 360.

Pnevmatikakis, E.A., Soudry, D., Gao, Y., Machado, T.A., Merel, J., Pfau, D., Reardon, T., Mu, Y., Lacefield, C., Yang, W., et al. (2016). Simultaneous Denoising, Deconvolution, and Demixing of Calcium Imaging Data. Neuron 89, 285–299.

Sun, F., Zeng, J., Jing, M., Zhou, J., Feng, J., Owen, S.F., Luo, Y., Li, F., Wang, H., Yamaguchi, T., et al. (2018). A Genetically Encoded Fluorescent Sensor Enables Rapid and Specific Detection of Dopamine in Flies, Fish, and Mice. Cell 174, 481–496 e419.

Zhou, P., Resendez, S.L., Rodriguez-Romaguera, J., Jimenez, J.C., Neufeld, S.Q., Giovannucci, A., Friedrich, J., Pnevmatikakis, E.A., Stuber, G.D., Hen, R., et al. (2018). Efficient and accurate extraction of in vivo calcium signals from microendoscopic video data. Elife 7.

